# Evolutionary context-integrated deep sequence modeling for protein engineering

**DOI:** 10.1101/2020.01.16.908509

**Authors:** Yunan Luo, Lam Vo, Hantian Ding, Yufeng Su, Yang Liu, Wesley Wei Qian, Huimin Zhao, Jian Peng

## Abstract

Protein engineering seeks to design proteins with improved or novel functions. Compared to rational design and directed evolution approaches, machine learning-guided approaches traverse the fitness landscape more effectively and hold the promise for accelerating engineering and reducing the experimental cost and effort. A critical challenge here is whether we are capable of predicting the function or fitness of unseen protein variants. By learning from the sequence and large-scale screening data of characterized variants, machine learning models predict functional fitness of sequences and prioritize new variants that are very likely to demonstrate enhanced functional properties, thereby guiding and accelerating rational design and directed evolution. While existing generative models and language models have been developed to predict the effects of mutation and assist protein engineering, the accuracy of these models is limited due to their unsupervised nature of the general sequence contexts they captured that is not specific to the protein being engineered. In this work, we propose ECNet, a deep-learning algorithm to exploit evolutionary contexts to predict functional fitness for protein engineering. Our method integrated local evolutionary context from homologous sequences that explicitly model residue-residue epistasis for the protein of interest, as well as the global evolutionary context that encodes rich semantic and structural features from the enormous protein sequence universe. This biologically motivated sequence modeling approach enables accurate mapping from sequence to function and provides generalization from low-order mutants to higher-orders. Through extensive benchmark experiments, we showed that our method outperforms existing methods on ∼50 deep mutagenesis scanning and random mutagenesis datasets, demonstrating its potential of guiding and expediting protein engineering.

## Introduction

The goal of protein engineering is to discover protein sequences that have a desired or enhanced property. However, the protein sequence space is tremendously large, and thus an exhaustive exploration or enumeration of the space is intractable in the laboratory and computationally when the length of the protein sequence is longer than 50 residues [1]. Furthermore, functional proteins are extremely rare in the astronomical sequence space, making it very challenging to successfully engineer a protein by a naive approach. Mainstream protein engineering strategies include rational design and directed evolution. Rational design optimizes the protein properties by making precise site-specific mutations based on detailed knowledge of structure, mechanism, and dynamics [2]. Rational design is oftentimes assisted by quantitative approaches to evaluate sequences that fold into a specific structure, including biophysical modeling [3,4], force field approximations [5–7], structure prediction [8–10], and molecular dynamics simulations [11]. Directed evolution is a protein engineering technique that circumvents this intractability in a greedy way [12–14] without the availability of protein structure. In directed evolution, a set of variants of the wild type sequence is constructed and then screened for the desired function. Variants with improved properties are selected to parent the next round of diversification until the desired functional fitness is achieved [1]. While having been demonstrated successful, the fraction of sequences that can be sampled by the most high-throughput screening and selection method is still very limited, and implementing an effective screen and selection can require a significant experimental effort [15].

To reduce the experimental effort, machine learning (ML) methods have been proposed to assist directed evolution and successfully engineered multiple important functional proteins [15–20]. In ML assisted evolution, a machine learning model is trained to learn the sequence-function relationship from sequence and screening data. In one round of evolution, the model simulates and predicts the fitness of all possible sequences, and a restricted list of best-performing variants are used as the starting point of the next round of evolution. In contrast to the classical directed evolution, ML assisted evolution is able to escape from the local optimum by learning the entire functional landscape from data. It takes full advantage of all available sequence and screening data, including those of unimproved variants, thereby traversing the fitness landscape more efficiently.

A critical component of ML guided directed evolution is to build a machine learning model that accurately maps sequence to function. Unlike the qualitative predictions that group protein sequences into different functional classes [21–23], in protein engineering, a model is required to distinguish quantitative functional levels of closely related sequences -- as in one round of directed evolution, the ML model needs to predict the fitness of a sequence that differs the parent sequence by only one or very few single amino acids. Several existing computational methods predict the mutation effects by leveraging the evolutionary information of homologous sequences [24,25]. Natural protein sequences we observe today are the result of a long-lasting natural evolution that selects proteins with desired functionalities. Following this intuition, these methods built generative models to reveal the underlying constraints of the evolutionary process, which can then be used to infer which mutations are more tolerable or favorable than others. Because of the unsupervised nature, however, these methods are not able to leverage the fitness data of tested variants available during the directed evolution process and thus may have limited accuracy when guiding the protein engineering. More recently, a trend is emerging that pre-trains a language model (LM) on large protein sequence datasets to learn protein sequence representation [26–30]. By trained to predict the next amino acid given all its preceding ones, the language model learns representations that are semantically rich and encode structure, evolutionary and biophysical contexts [27]. It was found that using the learned representation as the feature input to fine-tune a supervised model improves fitness prediction on multiple protein mutagenesis datasets [28]. However, as these models are trained on massive sequences such as those in UniProt [31] and Pfam [32], the learned representations only capture general context for a wide spectrum of proteins but may not be specific to the protein to be engineered. Lacking this specificity in the representation, the prediction model may not be effective in capturing the underlying mechanism (e.g., epistasis between residues) that determines the fitness of a protein and is not able to effectively prioritize best-performing variants to assist the directed evolution.

Here, we developed ECNet, a deep learning model that guides protein engineering by predicting protein fitness from the sequence. We constructed a sequence representation that incorporated the local evolutionary context specific to the protein to be engineered. This representation explicitly encodes the residue interdependencies of all residue pairs in the sequence, which informs our prediction model to quantify the effects of mutations -- especially higher-order mutations -- in the sequence. We further incorporated with global evolutionary context from an LM model trained on large sequence databases to model the semantic grammar within protein sequences as well as other structure and stability relevant contexts. Finally, a recurrent neural network model, trained on the fitness data of screened variants, is used for the sequence-to-function modeling with both representations. Through extensive benchmark experiments, we showed that our method outperforms existing methods on ∼50 deep mutagenesis datasets. Further experiments on combinatorial mutagenesis datasets demonstrated that ECNet enables generalization from low-order mutants to higher-orders.

## Results

### Residue co-evolution correlates protein functional fitness

Mutations within the protein sequence can affect fitness in a non-independent way, which is also known as genetic interactions or epistasis. It was found that epistasis interactions, quantified by deep mutagenesis scanning (DMS) of proteins, can be used to infer protein contacts and structures [33,34]. As structurally proximal protein residues are often inferred from co-variation pairs from sequence evolution historically [35,36], we hypothesized that co-evolution information can also be used to infer epistasis or fitness of proteins.

We investigated the relationship between the co-evolution of residue pairs and the fitness of double mutants. We collected a DMS study that measured the fitness of double mutants of the human YAP65 WW domain [37]. We also quantified the strength of pairwise residue dependencies by fitting a direct coupling analysis model [38] to the homologous sequences of the WW domain. We found that the strength of pairwise dependencies correlated with the fitness of double mutants (Spearman correlation 0.35; **Figure 1a**). Similar to a previous study [24], we also used the change of dependency strength (by contrasting the mutant sequence to the wild type sequence) to predict the fitness of protein variants in a set of DMS studies [39]. We found that the predictions correlated with experimental data with a Spearman correlation ranging from 0.1 to 0.5 (**Figure 1b**). In addition, we observed a trend of increasing correlation score if a protein has more homologous sequences, presumably because abundant homologous sequences lead to a more accurately fitted direct coupling analysis model. Overall, these results suggested that there are signals in the evolution information that we can leverage to predict protein fitness. This motivated us to integrate evolutionary information of protein sequences to empower a supervised model that predicts the fitness of protein variants in directed evolution.

**Figure 1.**
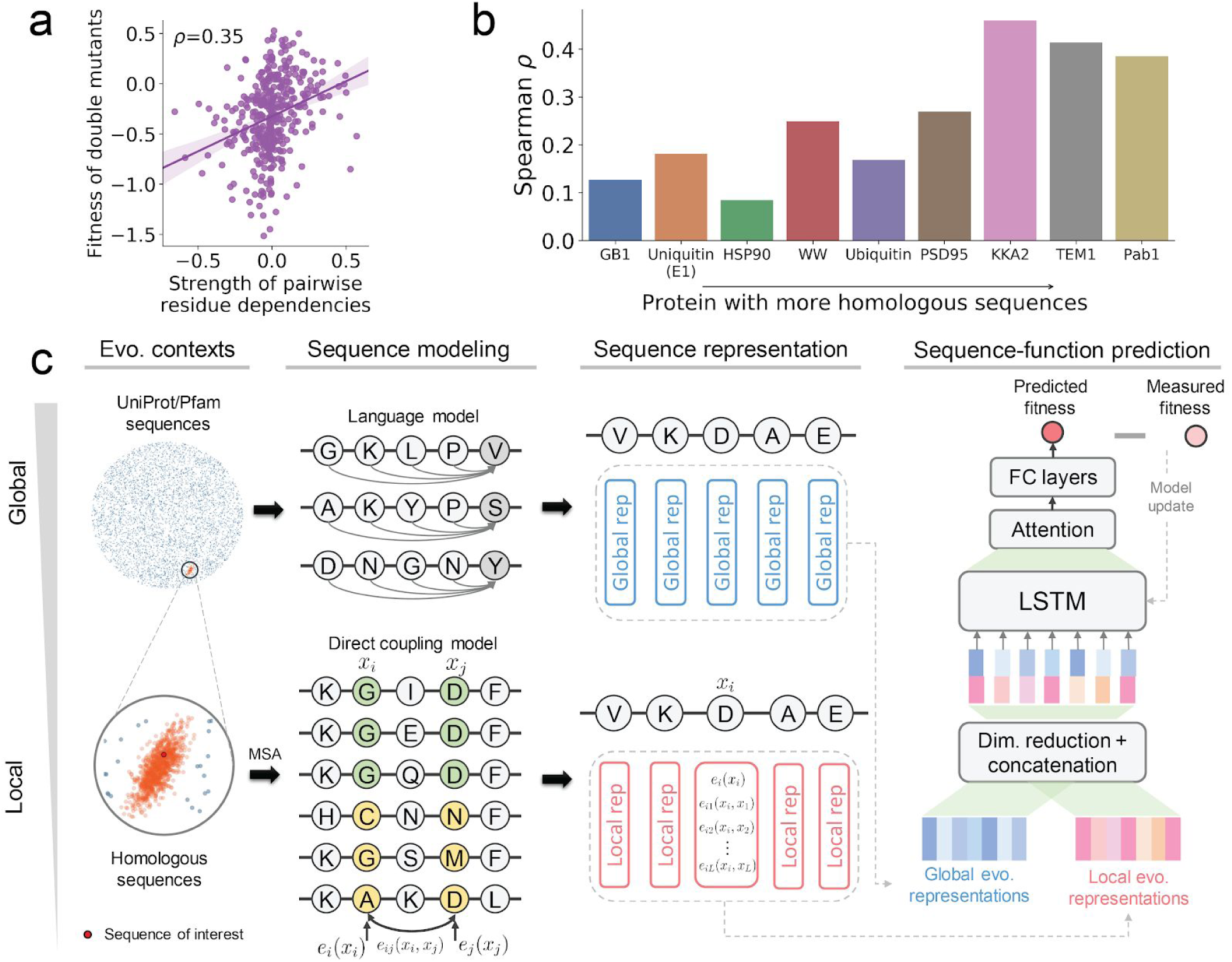
The motivation and overview of our evolutionary context-integrated sequence modeling method for protein engineering. **(a)** Sequence co-evolution data correlates with fitness measurements in deep mutagenesis scanning studies. The scatter plot shows the relationship between fitness measurement of double mutants and the co-variation strength of residues where the mutations were introduced. Each data point represents a double mutant. **(b)** Sequence co-evolution data can be used to predict protein fitness. The bar plot shows the Spearman correlation between experimentally measured fitness and strength changes of co-variation. Proteins were sorted by the number of homologous sequences. **(c)** An overview of our evolutionary context-integrated deep sequence modeling approach for protein engineering. Our method, ECNet, integrates global and local evolutionary contexts to represent the protein sequence of interest. First, a language model is used to learn global semantic-rich global sequence representations from the protein sequence databases such as UniProt or Pfam. Next, a direct coupling analysis model is used to capture the dependencies between residues in protein sequences, which encodes the local evolutionary context. The global and local evolutionary representations are then combined as sequence representations and used as the input of a deep learning model that predicts the fitness of proteins. Quantitative fitness data measured by deep mutagenesis scans (DMS) are used to supervise the training of the deep learning model. (MSA: multiple sequence alignment; Dim. reduction: dimensionality reduction; LSTM: long short-term memory network; FC layers: fully-connected layers; Evo. contexts: evolutionary contexts; Evo. representations: evolutionary representations.)

### Sequence-to-function modeling

We built a deep learning sequence-to-function model, ECNet, that learns the mapping from protein sequences to their respective functional measurements (**Figure 1c**) from data (e.g., fitness measured by deep mutagenesis scans). We used the LSTM neural network architecture and trained protein-specific models using large-scale deep mutagenesis scans datasets (**Supplementary Note**).

Our model is mainly empowered by two informative protein representations, with one accounting for residue interdependencies of the specific protein of interest and the other capturing the general sequence semantics in the protein universe. Existing tools predict the conservation effects of mutations by considering each amino acid independently (e.g., PolyPhen-2 [40] and CADD [41]) while others exploit structure information (e.g., FoldX [42] and OSPREY [4]). However, the functions of proteins are often driven by the interdependencies between residues (e.g., epistasis) in the protein [43,44], and not all the protein structures are solved. We thus explicitly modeled the pairwise interactions of all pairs of sites in a protein by extracting signals of evolutionary conservation from its homologous sequences or sequence families. We used a generative graphical model, fitted on the multiple sequence alignment (MSA) of the homologous sequences, to uncover the underlying constraints or interdependencies that defines the family of homologous sequences. These constraints are the results of the evolutionary process under natural selection and may reveal clues on which mutations are more tolerable or favorable than others. The generative model generates a sequence *x* with probability *p*(*x*) = *exp*[*E*(*x*)]/*Z*, where *E*(*x*) is the ‘energy function’ of sequence *x* in the generative model and *Z* is a normalization constant. We applied CCMpred, which is based on a Markov random field (MRF) specification to model the residue dependencies in protein sequences and the energy function of sequence *x* is defined as the sum of all pairwise coupling constraints *e*_*ij*_ and single-site constraints *e*_*i*_, where *i* and *j* are position indices along the protein sequence,

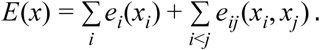

When the MRF model fit to data with proper regularizations, the residue interactions in protein sequences are explained by the direct coupling terms *e*_*ij*_. It has been shown that the magnitudes of *e*_*ij*_ terms can accurately predict protein contacts [10] and 3D structures [45]. For a protein sequence with length *L*, we encoded its *i*-th amino acid *x*_*i*_ by a vector, in which elements were set to the single-site term *e*_*i*_ (*x*_*i*_) and pairwise coupling terms *e*_*ij*_ (*x*_*i*_, *x*_*j*_) for *j* = 1, …, *L* (**Figure 1c**), and then dimensionality reduction techniques were used to project it into low rank (**Supplementary Note**). Encoding the protein sequence in this way directly incorporates the protein’s evolutionary context, i.e., the effects of pairwise epistasis, which can inform machine learning models to predict the fitness of a sequence with single or higher-order combinatorial mutations.

In addition to the evolutionary sequence contexts specific to the protein of interest, global protein sequence contexts, i.e., those encode structures and stabilities, can also inform our prediction model to predict the effects of mutations. For this purpose, we integrated general protein sequence representations from unsupervised protein contextual language models [26–29]. Using large corpora of protein sequences such as UniProt and Pfam, a language model learns to predict the next amino acid given all its preceding amino acids. During the training, the language model gradually changes its internal dynamics (encoded as hidden state vectors) to maximize the prediction accuracy. It was found that a wide range of protein relevant scientific tasks, including secondary structure prediction, contact prediction, remote homology detection, can be improved by using the hidden state vectors of a language model as input features to fine-tune a supervised model for the specific task [28,29]. Here, we also used the language model’s hidden state vectors as another type of protein sequence representation for our prediction model to capture the global protein sequence context (**Figure 1c, Supplementary Note**), which is a complement to our local evolutionary context representation.

The local and global evolutionary representations are jointly used to model the protein sequence of interest. A deep learning model (recurrent neural network) then takes these sequence representations as input and learns the sequence-to-function relationship. Quantitative functional measurements (e.g., fitness data measured by deep mutagenesis scans) are used to supervise the training of the deep learning model.

### Accurate prediction of functional fitness landscape of proteins

We performed multiple benchmarking experiments to assess the ability of ECNet in predicting the functional fitness from protein sequences.

We first compared our evolutionary context representation to different representation schemes for protein sequences or mutations. Yang et al [46] proposed to use a Doc2Vec model [47], pre-trained on ∼500k UniProt sequences, to map an arbitrary sequence to a 64-dimensional real-valued vector. To directly test the utility of sequence representations, we used our deep learning model as the predictor for both our representation and the Doc2Vec representation of Yang et al. We compared the two approaches on the Envision dataset [39], composed of 12 DMS studies that generated fitness values of single amino acid variants of ten proteins (**Supplementary Note**). We found that ECNet consistently outperformed the approach of Yang et al on all the 12 datasets, with a relative improvement ranging from 16%-60% (**Fig. 2a**). Since the Doc2Vec representation was learned from the UniProt dataset, the information it captured is mostly general protein properties but not the dependencies in the sequence that determine functions. In contrast, our evolutionary context representation explicitly models the epistasis of residue pairs in the sequence, which jointly influence the function in a non-independent way. This fine-grained information informed the prediction model to learn the sequence-function mapping more effectively and thus improved the prediction performance. We also compared our evolutionary context representation to the approach proposed in the Envision dataset paper [39], which described a single amino acid substitution using 27 biological, structural, and physicochemical features. Compared to this approach, our method, without using these features, still improved the Spearman correlation for 11/12 proteins (**Figure S1**). As protein engineering focuses on identifying variants with improved properties than the wild type, we further evaluated the model performance using a classification metric (AUROC score), in which variants with higher function measurements than the wild type sequence were defined as positive samples and the remaining variants as negative samples. We observed similar improvements in AUROC scores (**Figure 2b**). These results suggested that sequence contexts are more informative than the descriptors of mutated amino acids, which is critical in capturing the interdependencies between residues to predict the functions.

**Figure 2.**
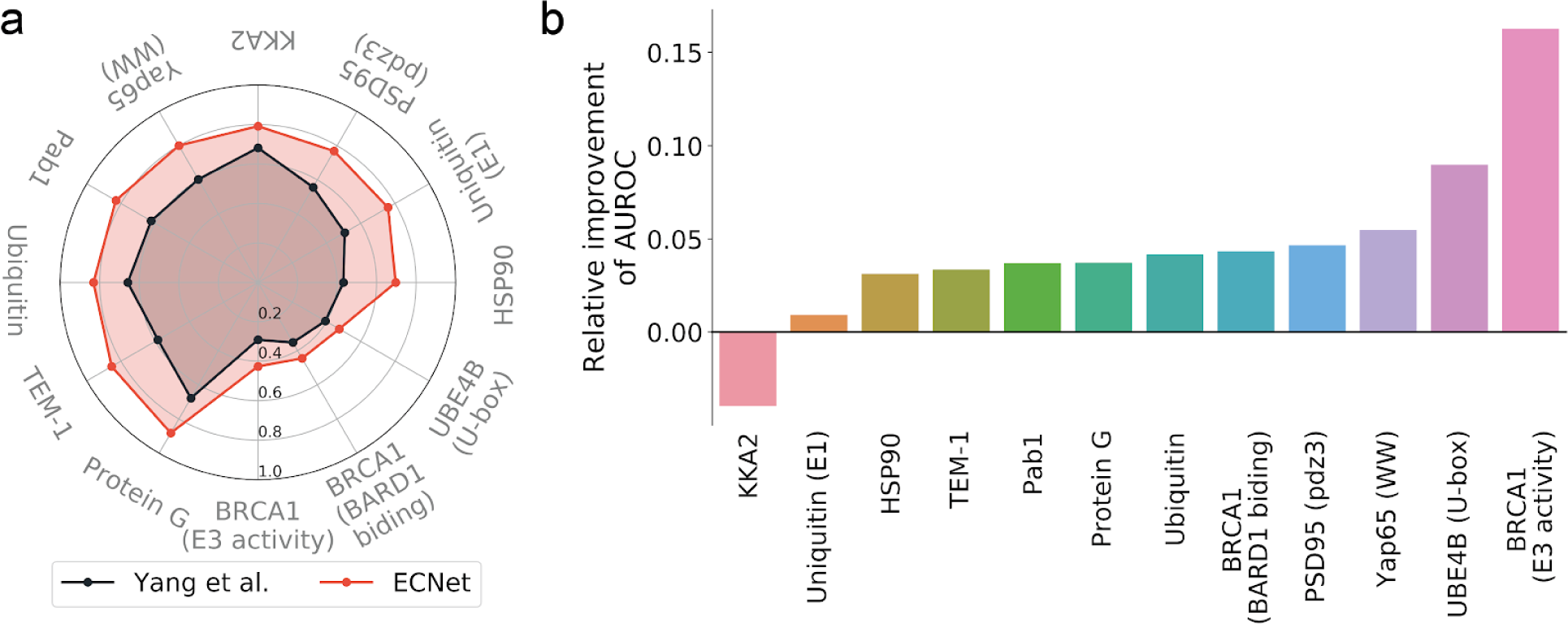
Comparisons to other protein variant representation methods. **(a)** Comparison to the approach from Yang et al. [46] that represents protein sequences with fixed-length vector representations by training a Doc2Vec model on the UniProt database. Spearman correlation was used as the evaluation metric. **(b)** Comparison to the Envision model [39] that represents a variant with 27 biological, structural, and physicochemical descriptors. AUROC was used as the evaluation metric to assess the ability of the model in identifying variants with improved function compared to the wild type. Relative improvements achieved by our method over the Envision model were shown in the barplot. All performances were evaluated using five-fold cross-validation.

Next, we compared ECNet to other sequence modeling approaches for mutation effects prediction on a larger set of DMS datasets previously curated in [25]. We first compared to three unsupervised methods, including EVmutation [24], DeepSequence [25], and Autoregressive [48]. These methods trained generative models on homologous sequences and predicted the mutation effects by calculating the log-ratio of sequence probabilities of mutant and wild type sequences. As expected, our method, predicting the mutation effects using a supervised predictor, outperformed these methods across almost all of the proteins (**Figure 3a**), compared to both EVmutation (median difference in Spearman correlation Δ`ρ`=0.193), DeepSequence (median Δ`ρ`=0.177), and Autoregressive (median Δ`ρ`=0.127). There were only two proteins on which our method did not clearly outperform other unsupervised methods. This is likely due to the relatively small number of function measurements available that we can use to train the supervised predictor (1,777 and 985 measurements, respectively; median: 2,721 across all proteins). We expected that a more regularized prediction model will achieve improved prediction performance for proteins with a small set of function measurements, which we leave as a future work. We also compared ECNet to a strong supervised baseline, in which the sequence representations learned by a language model were used as input to train a neural network predictor (denoted as Supervised LM). We used the architecture of the neural network of both our method and Supervised LM for fair comparison (**Supplementary Note**). We found that our method, by combining global LM representations, local evolutionary representations and the raw sequence as input, achieved higher correlations than the model that used LM representations alone for nearly all proteins (**Figure 3b**). We also performed an ablation analysis to dissect the performance of each representation component in our model’s input and found that a model used joint representations outperformed a model used any individual representation (**Figure S2**). Overall, tested on a large set of DMS data, our method significantly outperformed other sequence modeling methods, either unsupervised or supervised (**Figure 3c**; one-sided rank-sum test *P*<10^−5^), demonstrating its superior ability in predicting the fitness landscape of protein variants.

**Figure 3.**
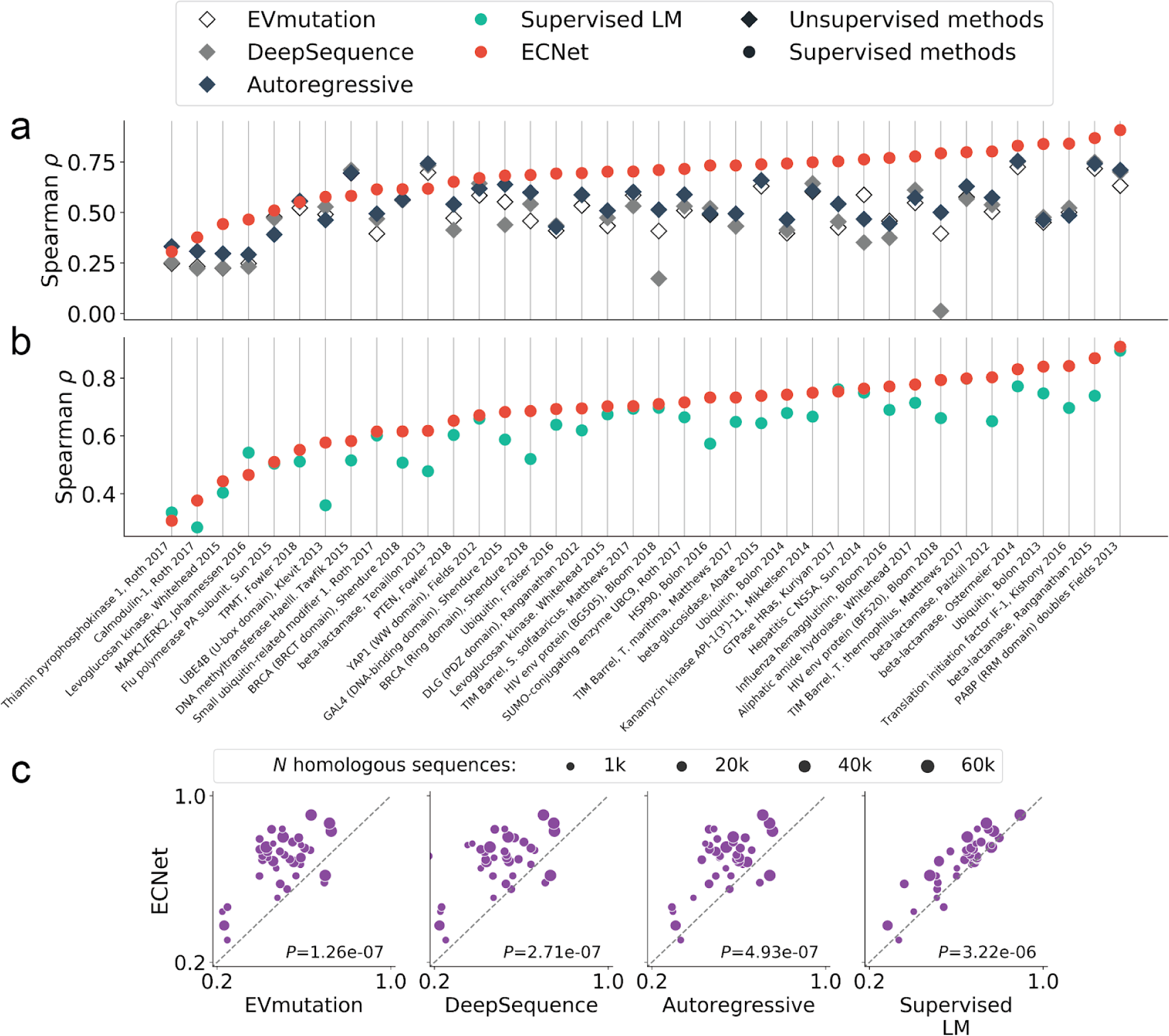
Comparisons to other sequence modeling approaches for mutation effects prediction. **(a)** Comparisons to three unsupervised generative models, EVmutation, DeepSequence, and Autoregressive. **(b)** Comparison to a supervised model that uses a pre-trained language model (LM) to learn protein sequence representation and fine-tunes a supervised predictor using functional measurements. **(c)** Pairwise comparisons between our method and other methods. Each data point represents the performance of one deep mutagenesis scans of a protein and the dot size is proportional to the number of homologous sequences of the protein. Spearman correlation was used as the evaluation metric for all results in this figure. One-sided rank-sum test was used to test the statistical significance.

### Generalization to higher-order variants from low-order data

The construction and screening of higher-order variants can require a significant experimental burden. As a result, fitness measurements of single mutants were more prevalent in previous DMS studies as compared to those of double or higher-order mutants. It is thus highly desired in protein engineering that a machine learning model trained on fitness data of low-order variants can also accurately predict the fitness of higher-order variants. As such, the model can fully leverage the fitness data of screened low-order variants and prioritize higher-order variants that are likely to exhibit improved properties for the next round of screen.

We thus assessed ECNet’s performance on predicting the fitness of higher-order variants when lower-order data were used for model training. We collected the fitness measurements of both single and double mutants of six proteins from previous DMS studies [37,49–53]. We then trained our prediction model using single mutants data only and tested its performance on double mutants. The model achieved Spearman correlation ranging from 0.73 to 0.94 for the six proteins and outperformed the Supervised LM and the EVmutation baselines (**Figure 4a**), suggesting its generalizability to the prediction of higher-order variants from low-order data. We also observed that increased diversity of fitness landscape in the training data improved the prediction performance. For example, to predict the fitness of quadruple mutants of the avGFP protein [54], we trained separate models using the fitness data of single, double, triple or all three orders of mutants. The test results suggested that a model trained on higher-order mutation data (from single to triple) achieved an increasing prediction performance, and the union of all-order mutation data further improved the prediction (**Figure 4b**). To further assess ECNet’s ability, we used orthogonal data containing sequences of 146 TEM alleles that are known to be inhibitor-resistant (**Supplementary Note**). Sequences in this data contain two to ten (mean 3.3) amino acid substitutions compared to the TEM protein. Based on these sequences, we generated ∼19k random variants by enumerating all mutation combinations restricted on the positions where mutations were introduced in the 146 alleles (**Supplementary Note**). We then trained our model on fitness data of TEM-1 single mutants data and used it to predict the fitness of the 146 TEM variants as well as the randomly generated variants. We found that our model distinguished the inhibitor-resistant variants from the random variant background (**Figure 4c**; mean predicted fitness 0.79 vs. 0.48; one-sided rank-sum test *P*<10^−5^). This orthogonal validation further demonstrated the generalizability of our model, even trained on single mutants data, to the prediction for higher-order mutants.

**Figure 4.**
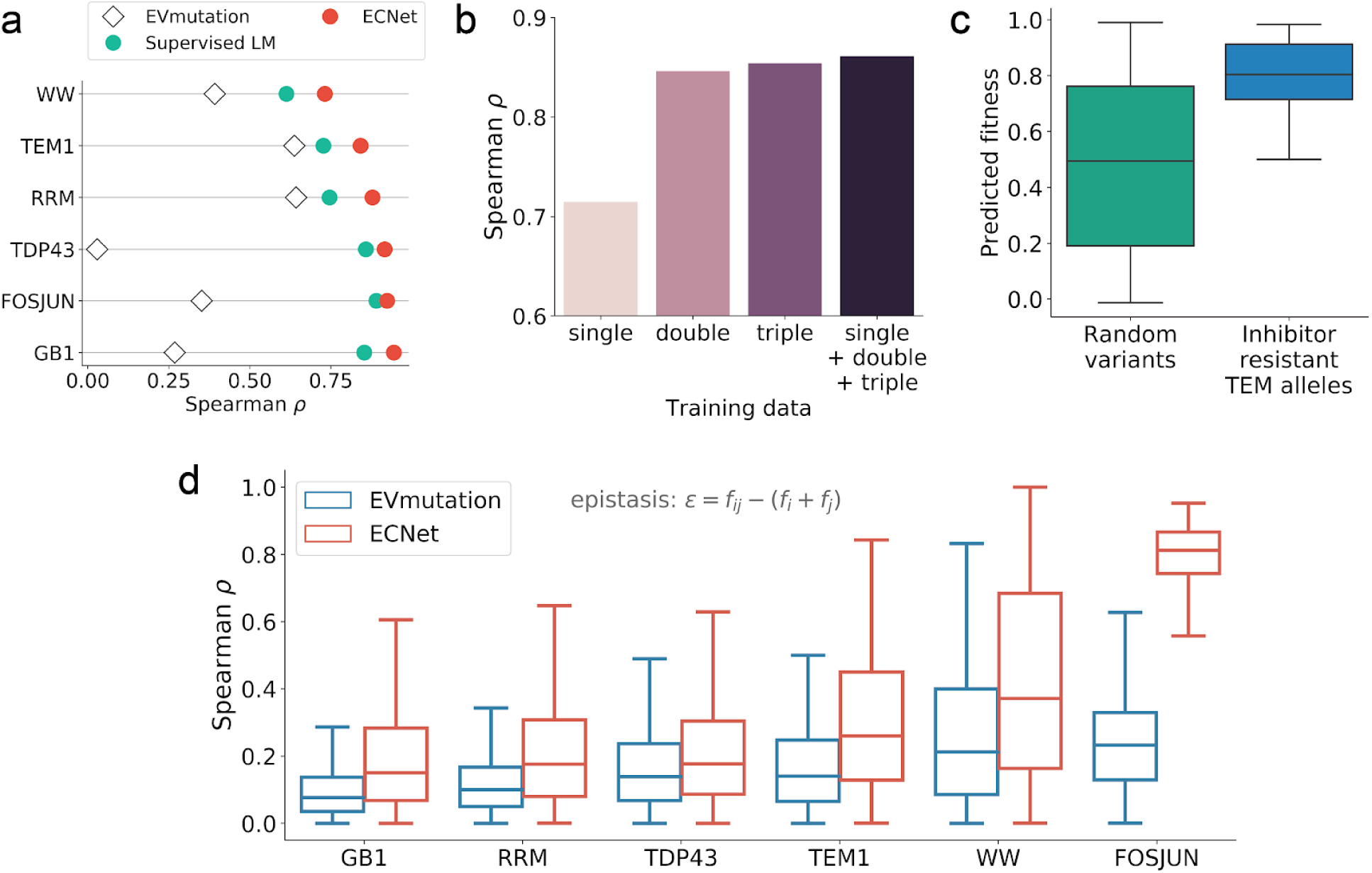
Accurate prediction of higher-order variants using a model trained on lower-order variants. **(a)** Prediction of the fitness of double mutants. For supervised methods (ECNet and Supervised LM), the prediction models were trained using fitness measurements of single mutants. **(b)** Prediction of quadruple mutants of avGFP using models trained on single, double, triple, and all three types of mutants. **(c)** The predicted fitness values of inhibitor-resistant TEM alleles were significantly higher than those of randomly generated background variants (one-sided rank-sum test *P*<10^−5^). **(d)** Spearman correlation of experimentally measured epistasis and predicted epistasis. The Spearman correlations achieved by our method were significantly higher than that of EVmutation for all six proteins (one-sided rank-sum test *P*<10^−4^).

Evidence has shown that mutations within the sequence can have non-independent effects (epistasis) on fitness [43,55]. The double mutant fitness *f* _*ij*_ may not always equal to the sum of constituent single mutant fitness *f* _*i*_ + *f* _*j*_, where *f*’s are the (log-transformed) experimentally measured fitness of variants. Epistasis (ε) is quantified as the difference between the experimentally measured fitness and the expected fitness: ε = *f* _*ij*_ − (*f* _*i*_ + *f* _*j*_). To analyze whether ECNet captures the interdependencies between mutations, we correlated the observed epistasis ε with predicted epistasis 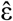, which is defined as 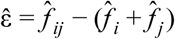 where 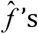 ‘s are predicted fitness. Compared with EVmutation that explicitly models epistasis using a generative model, the epistasis predicted by our method better correlated with the observed epistasis (**Figure 4d;** one-sided rank-sum test *P*<10^−4^). These results suggested that our method captured the residue dependencies within sequences more accurately, and thus resulted in the superior prediction performances reported above.

## Discussion

A critical challenge in machine learning guided protein engineering is the development of a machine learning model that accurately maps protein sequences to functions for unseen variants. While models have been developed for the qualitative classification of protein sequences into function classes, such as those in the Critical Assessment of Functional Annotation (CAFA) challenge [21], in protein engineering prediction models are required to provide a more fine-grained characterization of protein functions, which distinguishes the quantitative function levels of closely related sequences (e.g., single-site mutants of wild type protein with sequence similarity >99%). The function prediction in protein engineering is also different from predicting the deleteriousness [56] or instability [42] of variants -- to assist protein engineering, the machine learning model needs to prioritize variants that are not only structurally stable and non-deleterious but also with enhanced properties. Furthermore, as the protein sequence space is tremendous in size, it is desired to have a machine learning model that navigates the fitness landscape effectively and can generalize from regions of low-order variants to regions in the landscape where higher-order variants with improved function may exist. All these factors render it uniquely challenging to develop a machine learning model that can be used to guide protein engineering strategies such as directed evolution and rational design.

In this work, we have presented a high-performance method, ECNet, that predicts protein function levels from sequence to facilitate the process of protein engineering. The machine learning model used a biologically-motivated sequence modeling approach to learn the sequence-function relationship, leading to superior performances in predicting the fitness of protein variants. Benchmarked on a large set of deep mutagenesis scanning studies, the method outperformed a line of established methods. Further, our method accurately captured the epistasis effects of mutations within protein sequences and can be generalized to predict higher-order mutants’ functions by learning from the data of lower-orders.

We expect our method to be a practical tool for machine learning guided protein engineering. In a round of directed evolution, the sequence-to-function model can be applied to select the next set of variants to screen. In addition, given its generalizability to higher-order mutants from lower orders, the model can fully leverage the screening data of low-order mutants, including that of both improved and unimproved variants, generalize to distant regions in the fitness landscape where higher-order variants with improved properties may exist, and prioritize promising higher-order mutants to screen in the next round, in which the screened data can be used to further improve the model, hereby forming an iterative loop of directed evolution to discover improved variants.

## Acknowledgments

J.P. acknowledges the support from the Sloan Research Fellowship and the NSF CAREER Award. Y. Luo acknowledges the support from the CompGen Fellowship.

## Supplementary Information

### Supplementary Note

#### Datasets

We collected a set of large-scale deep mutagenesis scanning (DMS) datasets and a random mutagenesis dataset curated by previous publications.

##### Envision dataset

We first collected 12 DMS studies from [39], covering ten proteins and 28,545 fitness measurements of single amino acid variants. The fitness values were normalized such that wild type-like variants having scores of one, and variants that are more (less) activate than the wild type having scores greater (less) than one.

##### DeepSequence dataset

We also collected a set of DMS datasets compiled by [25]. We excluded a study of RNAs since it is out of the scope of this study. The resulting set consists of 38 DMS studies across 34 proteins. Most of these studies (36/38) provide the function values of single amino acid variants, and two studies provide the functional measurements of double or up to quadruple mutants. The functions measured in these studies include growth rate, enzyme function, protein stability, and peptide binding.

##### Single and double mutants datasets

To test the ability or our method to predict epistasis, we compiled multiple DMS studies that contain the fitness values of both single and double amino acid variants. We obtained the DMS data of the GB1 domain, WW domain, RRM domain, and FOS–JUN heterodimer from [34], and the prion-like domain of TDP-43 from [53]. A set of fitness of TEM-1 consecutive variants was also obtained from [52].

##### Higher-order avGFP mutants dataset

We also collected a higher-order mutants dataset [54] to assess our method’s generalizability to predict the effect of even higher-order variants. This study systematically assayed the local fitness landscape of the green fluorescent protein from Aequorea victoria (avGFP) by measuring the fluorescence of ∼50k derivatives of avGFP, with each sequence containing 1-15 amino acid substitution mutations.

##### Inhibitor-resistant TEM alleles

We compiled a list of TEM alleles that have been found to be inhibitor-resistant with supporting evidence in previous studies. The list was downloaded from https://externalwebapps.lahey.org/studies/TEMTable.aspx. We excluded alleles for which mutation information was labeled as “Not yet released”. This resulted in 146 sequences that mostly contains two to five and up to ten mutations (average 3.3 mutations per sequence). Based on this list, we also generated a list of random variants by enumerating all combinations of amino acid mutations on all or a subset of the positions where mutations were introduced in the 146-sequence list. In total, we obtained 18,937 randomly generated variants.

### Inference of evolutionary couplings from multiple sequence alignments

We first searched homologous protein sequences of a given protein using hhblits available in the hh-suite [57]. We used the wild type sequence of the given protein as the query sequence and searched against the uniclust-30 database (version uniclust30_2018_08) for three iterations. We used a maximum pairwise sequence identity of 99%. Other parameters were set as default. The search results were formatted to the A2M multiple sequence alignment (MSA) format.

To identify the co-evolutionary residue pairs in a protein, we used a statistical model to exploit the evolutionary sequence conservation and model all pairwise interdependencies of residues. The model identifies the evolutionary couplings by learning a generative model of the MSA of homologous sequences using a Markov random field. Given the MSA of homologous sequences, the couplings are learned by maximizing the likelihood of observed sequences in the MSA, which is defined as

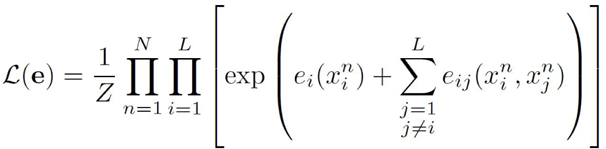

where 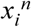 is the *i*-th amino acid in the *n*-th sequence, *Z* is the normalization constant, *N* is the number of homologous sequences and *L* is the number of columns in the MSA. The direct optimization of this likelihood is computationally intractable due to the computation of the normalization constant that increases exponentially -- 20^*L*^ sequences need to be considered. It was thus adopted to maximize the site-factored pseudo-likelihood of the MSA, which has a running time complexity *O*(*NL*^2^) where *N* is the number of sequences in the MSA. We refer the interested readers to [38,58,59] for the details of the optimization. In this work, we used CCMPred [38], a GPU based algorithm maximizing the pseudo-likelihood (plus regularization terms), to optimize the generative model. The evolutionary couplings are learned as parameters of the Markov random field.

### Local evolutionary context representation with evolutionary couplings

By fitting the graphical model to the MSA of homologous sequences of a protein, we obtained the coupling matrix *e*_*ij*_ that quantifies the co-constraints of all possible 20^2^ amino acid combinations between positions *i* and *j* in the sequence. In particular, the term *e*_*ij*_ (*x*_*i*_, *x*_*j*_) is the pairwise emission potential of the Markov random field for amino acid *x*_*i*_ occurring at position *i* while amino acid *x*_*j*_ occurring at position *j*. We use the coupling matrix *e*_*ij*_ to construct a data representation that encodes the co-evolution information of a protein.

Specifically, the *i*-th amino acid *x*_*j*_ in the protein was represented by an (*L+1)*-long ‘local evolutionary representation’:

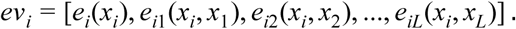

The full representation of a protein sequence was thus obtained by stacking local evolutionary representations for all positions, resulting in an *L* by *(L+1)* matrix. As we have shown, the pairwise potentials in matrix *e*_*ij*_ correlated with the fitness measured in DMS experiments (**Figure 1a-b**). We thus expect that using the local evolutionary representations derived from *e*_*i*_ and *e*_*ij*_ as a data representation of amino acids will inform a prediction model (described below) to better capture the residue dependencies and the sequence-to-function relationship.

The length of the local evolutionary representation is roughly equal to the length of the protein sequence, which may raise an overfitting issue when the protein length is long while the number of functional measurements used as training data is low. Therefore, we used a dimensionality reduction approach to transform the *(L+1)*-long vector into a fixed-length *d*-dimensional vector, where *d* is independent with the length of the protein sequence. To do this, we first encoded the protein sequence of each variant (with amino acid substitutions with respect to the wild type sequence) in the MSA. Next, we applied a position-specific PCA to the local evolutionary representation *ev*_*i*_ of position *i* and kept the first *d* principal components as the transformed local evolutionary representation. We set *d*=32 in this work. In the model testing stage, the dimensionality reduction was performed to each position using the PCA transformation fitted on training data. Hereinafter, we will refer to the PCA-transformed vector *ev*_*i*_ as local evolutionary representation unless otherwise specified.

### Pre-trained protein sequence representation model

Very recently, self-supervised models have provided powerful protein sequence representations that facilitate scientific advances, including protein engineering, structure prediction, and remote homology detection. These language models [26–29], without using labeled data, are trained on natural sequences from large protein databases such as Pfam [32] and UniProt [31] to predict the next amino acid character given all previous amino acid characters in the protein sequence or predict randomly masked amino acids using the rest. During the model training, these models progressively adapt their parameters to maximize the prediction accuracy, resulting in a representation of protein sequences that capture intrinsic semantics in protein sequences and interdependencies among amino acids.

In this work, we integrated the amino acid representations by a transformer model in TAPE, one of the most powerful self-supervised sequence representation models [29]. The representations capture the global evolutionary context from the massive protein sequence data the model was trained on, which is complementary to our evolutionary representations that capture the local evolutionary context. The TAPE model applied a Transformer architecture [60] and was trained on Pfam data to predict a masked amino acid using the remaining ones. We downloaded the pre-trained weights of the TAPE model from https://github.com/songlab-cal/tape. For an input sequence, TAPE generates a 512-dimensional vector representation for each amino acid. To integrate the TAPE representations into our model, we reprojected them to 128-dimensional vectors using a linear fully-connected layer, which were then concatenated with the local evolutionary representations and one-hot encodings and passed to the LSTM model. We refer to the reprojected TAPE representations as global evolutionary representations.

### Sequence-to-function neural network model

#### Model architecture

We built a deep learning model for the sequence-to-function prediction. The model receives as input features (amino acid characters and evolutionary representations) of the protein sequences and produces the predicted functional measurements of proteins as output. The backbone of our model is a long short-term memory network (LSTM) [61] integrated with a two-layer fully-connected neural network. Amino acids in the input sequences were one-hot encoded and passed through a 20-dimensional embedding layer. The amino acid embeddings were then concatenated with the evolutionary representations position-wisely before being input to the LSTM module. We used a single-layer LSTM with a hidden dimension *d*_*LSTM*_ = 128 as the default setting in this work. One hidden state vector was produced by the LSTM for every amino acid in the sequence. We summarized these hidden state vectors into a single vector using a weighted average, where the averaging weights were learned from the data by using a self-attention layer [60]. This vector was then passed to a top module to predict the functional measurements. The top module is a two-layer fully-connected neural network with tanh activation. The hidden dimensions of the two layers were set to 64 and 1, respectively. To facilitate the model training, we added a batch normalization layer [62] before the fully-connected layers. We also applied a dropout [63] layer after the first fully-connected layer to prevent overfitting.

#### Training details

We cast the task of predicting the functional values of proteins as a regression problem, and the objective was to minimize the difference between the predicted and experimentally measured functional values. We trained our deep learning model using the Adam optimizer [64] with default parameters. Mean squared error (squared *L*_*2*_ norm) was used as the loss function. The batch size was set to 128 and the maximum number of training epochs was set to 1,000 with an early stop if the performance has not been improved for 500 epochs. Model training was performed on an Nvidia TITAN X GPU. The training time depends on the training data size of each protein, ranging from 0.5 to 6 hours.

#### Auxiliary classification objective

While the prediction functional measurements is a regression problem by definition, the skewed distribution of the training data may lead to a biased predictor. For example, in the Envision dataset [39], only 18% of TEM-1 variants are more active than the wild type sequence (positive effects) while the remaining are less active than the wild type sequence (negative effects). In this case, a model optimized using a regression objective (e.g., minimizing the mean squared error) tends to fit the negative effects more but be less sensitive to the error from the prediction of positive effects. However, the main goal of machine learning guided protein engineering is to identify the variants with a better property than the wild type sequence. Hence, it is critical to mitigating this type of bias in the prediction model. We addressed this issue by introducing an auxiliary classification objective. We binned the functional measurements using their 10-quantiles as breakpoints, i.e., grouping the measurements into 10 bins with equal size. In the model training, we encouraged the model to accurately predict not only the absolute functional measurement but also which bin the measurement is in. Jointly, the classification objective forces the model to treat each interval of functional measurements equally and the regression objective forces the model to predict the measurements as close to the observed values as possible. Specifically, we added a second top module into the deep learning model, which also receives the summarized LSTM hidden state vector as input and its output are ten numbers indicating the predicted probability that the measurement should fall in each of the bins. The overall loss function is *L* = *L*_*rgr*_ + α*L*_*cls*_ where *L*_*rgr*_ is the loss of the regression objective, *L*_*cls*_ is the loss of the classification objective, and α set as α = 0.1 in this work. is a constant used to balance the scales of the two losses, which was

### Baseline methods

We compared our method against several existing baseline methods, including supervised and unsupervised models.

#### Yang et al. (Doc2Vec)

Yang et al. [46] proposed a learned embedding protein embedding to represent a protein sequence in a 64-dimensional vector using a Doc2Vec model [47] trained on the UniProt database. The representation vector is used as the input feature to fit a Gaussian process based regressor to predict the functional measurement. Following a previous work [27], we also used the four best performing models as chosen in Yang et al. [46], including the original model (k=3, w=7), the scrambled model (k=3, w=5), the random model (k=3, w=7), and the uniform model (k=4, w=1). The pre-trained models were downloaded from http://cheme.caltech.edu/~kkyang/models/ and protein representation vectors were generated using the code available at https://github.com/fhalab/embeddings_reproduction. The best performance across the four models was reported as the final performance of the Doc2Vec model.

#### Envision

Envision is a supervised method proposed in [39] that predicts the functional measurements of protein variants. Each variant was annotated with 27 biological, structural, and physicochemical features, which were used as input to train a gradient boosting regression model using large-scale mutagenesis data. We downloaded the source code of Envision from https://github.com/FowlerLab/Envision2017.

#### Evmutation

EVmutation is an unsupervised statistical model proposed by Hopf et al. [24]. It explicitly models the co-variations between all pairs of residues in the protein by fitting a pairwise undirected graphical model to the multiple sequence alignment (MSA) of all homologous sequences of the protein of interest. The model then quantifies the effect of single or high-order substitution mutations using the log-ratio of sequence probabilities between the mutant and wild type sequences.

#### DeepSequence

Similar to EVmutation, DeepSequence [25] is also a generative model that predicts the effects of mutations in an unsupervised manner. However, unlike EVmutation explicitly modeling pairwise dependencies, DeepSequence uses a latent model, fitted on the MSA of homologous sequences of a protein, to capture higher-order dependencies of residues in the protein. The effects of mutations are also predicted by the log-ratio of mutant likelihood to wild type likelihood.

#### Autoregressive model

Generative models of protein sequences such as EVmutation and DeepSequence are dependent on the alignment of homologous sequences, which may introduce artifacts and lose important information caused by indels in the alignment. A generative autoregressive model was proposed by Riesselman et al [48] to predict the mutation effects in protein sequence, without the requirement of multiple sequence alignment.

#### Supervised LM

We used TAPE [29], a language model (LM) trained on Pfam sequences to generate global context representations of protein sequences. We extracted the hidden state vectors, one for each amino acid in the sequence, from the TAPE model. We used the same top module as in our model (i.e., self-attention layer and fully-connected layers) to take the representations as input and predict the functional measurements.

### Benchmarking experiments

To assess our method’s performance, we compared our method to other baselines using the original benchmark datasets that these methods were tested on in their publications.

#### Benchmarks on the Envision dataset

We compared our method to the gradient boosting regression algorithm (denoted as ‘Envision’) proposed in the Envision dataset paper [39]. We used five-fold cross-validation to evaluate model performance and the average performance across the five folds was reported as the final performance of a method. Spearman correlation was used as the evaluation metric. Note that Envision used 27 biological, structural, and physicochemical features to build the prediction model while our model only used the protein sequence to predict the functional measurements. To test the model’s ability to identify variants that are more active than the wild type sequence, we also converted the task into a classification problem, in which protein sequences with a function score greater than the wild type sequence (with a function score 1) were labeled as positive samples, and the remaining sequences as negative samples. We used the AUROC score as the metric for this classification evaluation. We also compared our method to the Yang et al. (Doc2Vec) model on this dataset.

#### Benchmarks on the DeepSequence dataset

We compared our method to EVmutation, DeepSequence and the Autoregressive model on the DeepSequence dataset that these methods have been tested on. The predictions made by these unsupervised approaches were collected from [25,48]. For our method, we performed five-fold cross-validation on this dataset and report the average performance over all the five folds. We used Spearman correlation as the evaluation metric. We also compared our method to the global context representation based model (Supervised LM) on this dataset.

## Supplementary Figures

**Figure S1.**
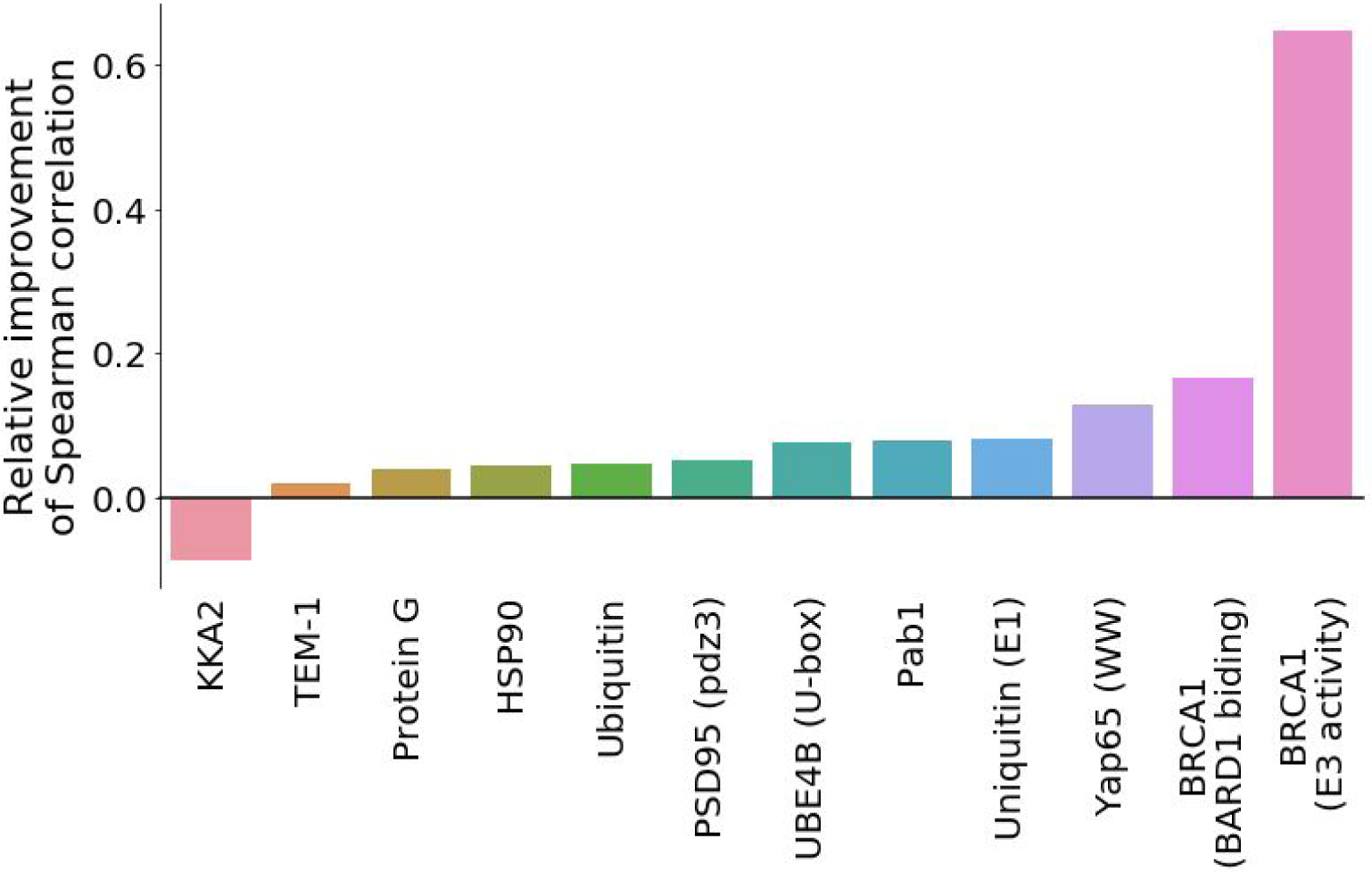
Performance improvements on the Envision dataset. This bar plot shows the relative improvements of Spearman correlation achieved by our method as compared to the Envision model. Related to **Figure 2b**.

**Figure S2.**
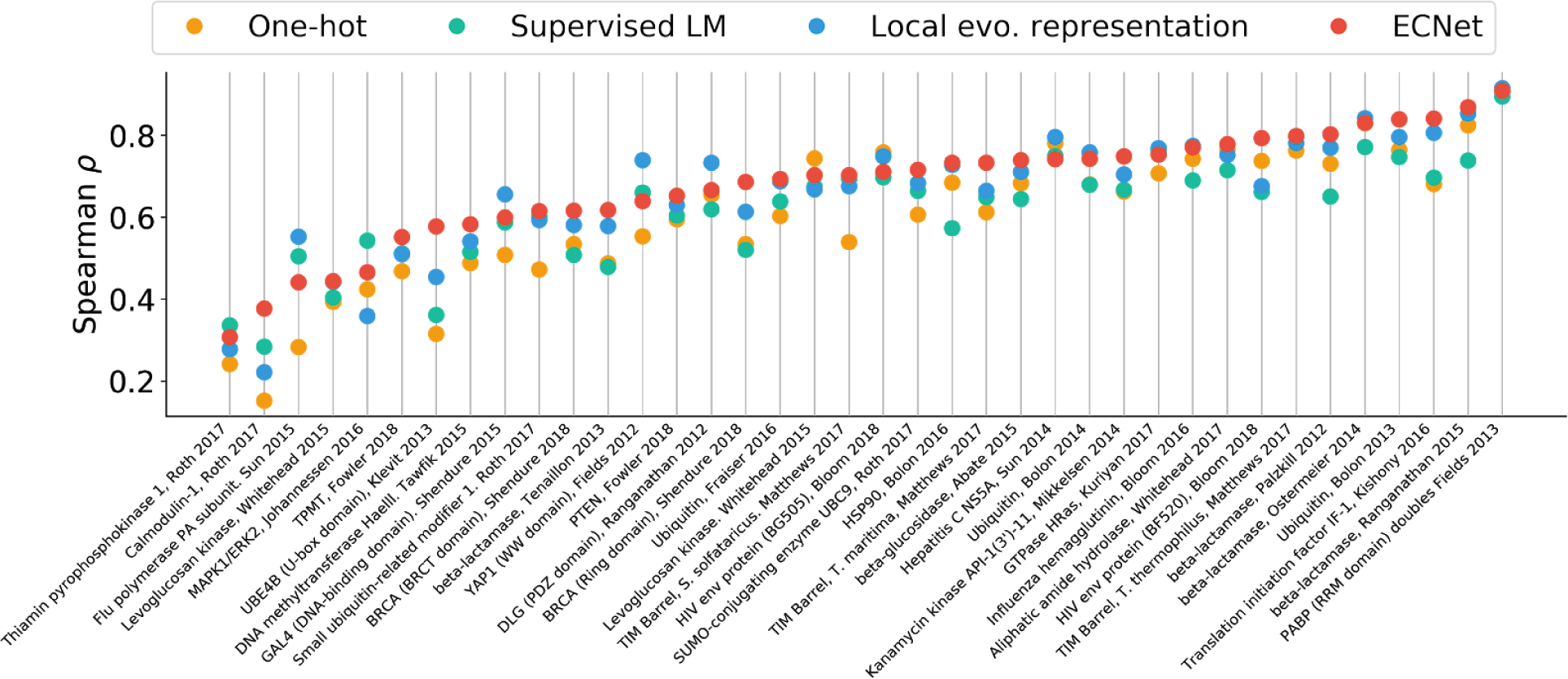
Comparison of supervised learning with different representations. ECNet integrated multi-scale sequence representations, including the global evolutionary representation, the local evolutionary representation and the one-hot representation of a protein sequence. We performed ablation analysis to compare ECNet to models that used individual sequence representation using five-fold cross-validation. Spearman correlation was used as the evaluation metric. Related to **Figure 3b**.

